# Panalyze: automated virus pangenome variation graph construction, analysis and annotation

**DOI:** 10.1101/2025.04.10.646565

**Authors:** Chandana Tennakoon, Thibaut Freville, Tim Downing

## Abstract

**Motivation:** Constructing and studying pangenome variation graphs (PVG) supports new insights into viral genomic diversity. This is because such pangenomes are less prone to reference bias, which affects mutation detection. Interpreting the information arising from this is challenging, so automating these processes to allow exploratory investigations for PVG optimisation is essential. Moreover, existing methods do not scale well to the smaller virus genome sizes and to facilitate analysis in laptop environments. To address this, we developed an easily deployable pipeline to facilitate the rapid creation of virus PVGs that applies a broad range of analyses to these PVGs.

**Results:** We present panalyze, a scalable and unbiased virus PVG construction, analysis and annotation tool implemented in NextFlow and containerised in Docker. Panalyze uses NextFlow to efficiently complete tasks across multiple compute nodes and in diverse high-performance computing environments. Panalyze can also operate on a single thread on a standard laptop, and analyse sequences lengths are of any size. We illustrate how Panalyze works and the valuable outputs it can generate using a range of common viral pathogens.

**Availability:** Panalyze is released under a MIT open-source license, available on GitHub with documentation accessible at https://github.com/downingtim/Panalyze/.

**Supplementary information:** Supplementary data are available online.

## 1. Introduction

Pangenome variation graphs (PVGs) offer new insights into genome evolution over short and long scales. A PVG is a sequence graph whose nodes represent sequences and whose edges represent connections observed in data (Hein 1989) such that a genome can be decoded by traversing a series of paths (Garrison et al 2018). PVGs combat reference allele bias by allowing reads to map to the most similar combination of paths in a PVG, instead of a single haplotype (Paten et al 2018). Reference bias is not obviated by using a consensus reference sequence or by using a multi-sample reference (Chen et al 2021, Chen et al 2024). This affects the chance of a read mapping to a location, impacting mutation detection, phylogenetic analyses, and associated inferences in pathogens (Valiente-Mulor 2017) particularly at variable regions that are often strongly associated with phenotypic changes (Haga et al 2024). This results in unaligned reads, missed mutations, poorer inferences, and less precise evolutionary models (Baaijens et al 2017, Hickey et al 2020). Novel pathogen isolates genetically distinct from known references will be characterised less accurately (Boehm et al 2021, Garrison 2023). In viruses, although genome assembly can be effective, extreme AT/GC content, host/vector contamination, short output contig lengths, and repetitiveness all contribute to incomplete and biased genomes (Alser et al 2021, Lischer et al 2017). Moreover, contigs often need to be aligned to existing databases either as sequences or k-mers, or an existing unavoidably-biased scaffold may be used as a reference (Bradley et al 2019, Ondov et al 2016).

Little work on viral PVGs has been completed (Downing 2025): only one study has yet been published: in this, PVGs helped recover >8% of sequence that failed to align to known linear reference genomes (Duchen et al 2024). A PVG can efficiently represent the diversity of multiple viral strains and so act as a better reference structure than a single linear reference. This means reads can map to different parts of the haplotypes represented in the PVG, which reduces the effect of known mutations and improves mutation discovery (Paten et al 2017, Rakocevic et al 2018). This circumvents the issue of reference genetic distance, improves alignment quality, and may permit more accurate genetic reconstruction of the sample collection (Garrison et al 2023, Moshiri et al 2022). Thus, there is a need for PVG creation and analysis to explore parameter spaces and optimise processes for viral PVGs.

There are many tools available for PVG creation and analysis. Most of these are developed with eukaryotic genome sizes in mind. These may require minimum sequence lengths larger than a typical virus genome length. Moreover, many of these tools are hard to install due to complex installation procedures, assume numerous dependencies, have steps requiring root access. Thus, it would be beneficial to have a curated set of pre-installed tools that can be easily deployed to automate PVG construction, analysis and annotation for small genome sizes. An existing tool, nf-core/pangenome (Heumos et al 2024) achieves this for larger eukaryotic genome sizes, but does not work for genomes <5 Kb long, has limited analysis options, and requires a minimum number of computer server threads. However, the smaller genome sizes of viruses means that their PVG analysis is possible on a laptop.

To address these issues, we present Panalyze, a scalable and rapid virus PVG construction, analysis and annotation tool operating in NextFlow v24.10.3 (Di Tommaso et al 2017) and containerised in Docker (Merkel 2014). Panalyze wraps a series of tools to allow sequence retrieval, PVG creation, phylogenetic analysis, PVG visualisation using multiple methods, PVG size calculation, PVG feature summarisation using different tools, PVG openness analysis, VCF generation, BUSCO gene detection, PVG community detection and PVG annotation. Panalyze focuses on constructing PVGs for smaller (viral) genomes and on supporting the interpretation and analysis of PVGs. Panalyze can automatically: download data of interest, align it, make a phylogeny, compute additional PVG metrics, identify BUSCO genes, annotate the PVG and compare inter-sample genome coordinates. Panalyze can use cluster nodes efficiently in a scalable manner that can be implemented on a single thread on a laptop or using many threads on a computer server. It also resolves the issue of complex installations and dependencies using Docker. We illustrate how Panalyze can work effectively for segmented, DNA and RNA viruses.

## 2. Methods

### 2.1 Pipeline overview

Panalyze has two entry points: a pre-existing FASTA file, or a search query from the user (Figure 1A). The latter seeks all complete genomes or genomic sequences matching the query in the Nucleotide database (Sayers et al 2021). The sequences are aligned with Mafft v7.453 (Katoh & Standley 2013) using automatic optimisation and default parameters. Evolutionary relationships are reconstructed using RAxML-NG (Randomised Axelerated Maximum Likelihood) v1.2.0 (Kozlov et al 2019) with a GTR (general time reversible) model and gamma substitution rate heterogeneity. Other models can be selected with modeltest-ng (Darriba et al 2020). Phylogenies are mid-pointed rooted and visualised using R v4.3.2 (R Core Team 2025) packages ape v5.7-1 (Paradis & Schliep 2019), ggtree v3.8.2 (Yu et al 2017), phangorn v2.11.1 (Schliep 2011), Rcpp v1.0.11 (Eddelbuettel et al 2024b), RcppArmadillo v0.12.6.6.0 (Eddelbuettel et al 2024a), phytools v2.0-3 (Revell 2024) and treeio v1.24.3 (Wang et al 2020).

**Fig. 1.**
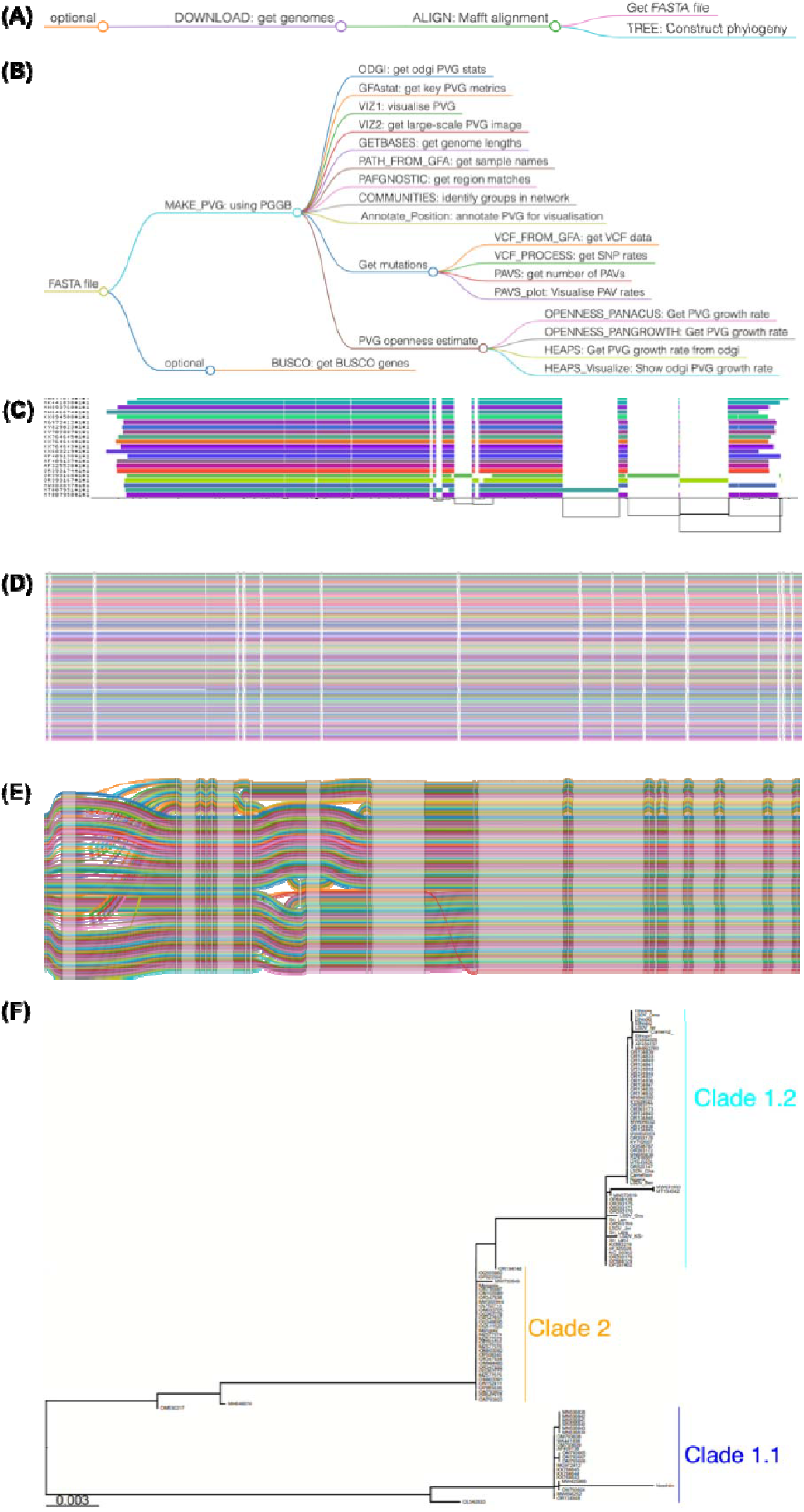
A representation of Panalyze’s workflow processes (right). Users can either (A) retrieve sequences of interest via the DOWNLOAD process to get FASTA input for (B), or curate a sample set and start the pipeline at (B). (B) MAKE_PVG will take the FASTA to make the PVG, before running analyses, including: getting PVG metrics with the ODGI and GFAstat processes, visualisations with VIZ1 and VIZ2, genome lengths with GETBASES, creating PAFs with PAFGNOSTIC, community detection with COMMUNITIES and annotation. Other processes are on mutation detection: determining VCFs with VCF_FROM_GFA, VCF_PROCESS to infer mutation rates, getting PAVS with PAVS, and visualising the PAVs with PAVS_plot. Users can implement openness analyses using Panacus, Pangrowth and ODGI, HEAPS and HEAPS_Vizualize. Users can (optionally) identify Busco genes with BUSCO. (C) A region at 135-140 Kb from a 121-sample LSDV PVG visualised using viz in ODGI where each genome shown is a coloured bar and the PVG topology is shown as the genomic coordinates the x-axis (black lines). (D) A region at 136-138 Kb of the same PVG visualised with viz in VG. (E) A region in the same PVG visualised using SequenceTubeMap, showing one minor (22 samples from Clade 1) and one major (99 samples from Clades 1.2 and 2) extended haplotype at LD144 (encoding a kelch-like protein at 135,541-137,208 bp) and LD145 (encoding an ankyrin repeat protein at 137,232-139,136). (F) A phylogeny based on this region showing the two main divergent haplotypes.

Panalyze creates PVGs using the pangenome graph builder (PGGB) v0.7.2-27 pipeline (Garrison et al 2024), which initiates all-to-all alignment with WFMASH v0.14.0-0 (Guarracino et al 2021) subject to a threshold of 90% identity per Kb (Figure 1B). PGGB keeps all paths and was used to reconstruct the input haplotypes directly. PGGB creates the PVG using SEQWISH (Garrison & Guarracino 2023) using a k-mer of 19 bp and a window size of 27 bp, and then sorts and orders it to produce a progressively linearised PVG based on partial order alignment with SMOOTHXG v0.7.0. This also removes erroneous redundant nodes (Garrison & Guarracino 2023). To allow analysis of a region < 1.5 Kb in size, we pad the sequences with nucleotides to a minimum length of 1.5 Kb. Once a PVG is built, we break the PVG into single-nucleotide nodes, retaining the original topology. Then we remove the nodes belonging to the padded sequences. Finally, we collapse this pruned PVG so that consecutive unitigs are collapsed into a contig. Multiqc v1.14 (Ewels et al 2016) collates PGGB metrics. The PVGs are generated in Graphical Fragment Assembly (GFA) format version 1. Panalyze does not use PVG pruning, chunking or lacing because virus PVGs are small. These PVGs are converted with the convert function in variation graph (VG) v1.43.0 (Garrison et al 2018), indexed with VG’s autoindex function. The corresponding FASTA file is indexed with SAMtools (Li 2009).

Next, a range of PVG analyses are applied. GFASTATS v1.3.6 (Formenti 2022), and ODGI (optimized dynamic genome/graph implementation) v0.8.3 (Guarracino et al 2022) produce PVG summary statistics. Visualisations are created with ODGI (Guarracino et al 2022) (Figure 1) and PGGB (Garrison et al 2023). The number of bases in the PVG comes from ODGI (Guarracino et al 2022). A VCF is produced by GFAUTIL v0.3.2 (Fischer 2021). The relative differences between the sequences’ coordinate systems is computed by GFAUTIL and visualised with ggplot2 v2_3.4.4 (Wickham 2016). PVG openness is computed firstly by Panacus v0.2.3 (Parmigiani et al 2024) using the nodes in the PVG with default coverage thresholds that included all segments, secondly with k-mers using Pangrowth (Parmigiani et al 2024), which also estimates the core PVG size, and thirdly with sequences extracted from PVG nodes using ODGI’s heaps function using 1,000 simulations (Guarracino et al 2022). Three methods are used to ensure consistency of openness estimates. PVG communities are created by PGGB (Garrison et al 2023) where the number of clusters can be determined by the user and the file names are indexed using fastix. This PVG community data is parsed, visualised and pairwise alignment format (PAF) files are generated for input to PAFGNOSTIC to numerically summarise PVG properties. A tabular PVG annotation file is generated using Prokka (Seeman 2014) for visualisation with Bandage (Wick et al 2015). Optional analyses include determining key gene presence-absence via BUSCO (Manni et al 2021). Additionally, the presence-absence variation (PAV) can be determined by ODGI (Guarracino et al 2022) and visualised, but should only be used for smaller or less diverse datasets due to the compute time required.

### 2.2 Testing the workflow

To illustrate that Panalyze’s PVG analysis approach was broadly applicable to all virus genomes, a range of test datasets were analysed (Table 1). Firstly, we applied it to a set of 121 lumpy skin disease virus (LSDV) genomes, representing large DNA viruses (taking 114 CPU hours, 78 minutes to complete). Secondly, we evaluated Panalyze’s performance on 15 porcine respiratory coronavirus (PRCV) genomes, representing small ssRNA viruses (6.7 CPU hours, 6.8 minutes). We repeated this on (ssRNA) 18 FMDV serotype C genomes (2.0 CPU hours, 2.6 minutes), and 441 FMDV serotype O genomes (353 CPU hours, 518 minutes). Thirdly, we evaluated its effectiveness in interpreting a segmented ssRNA virus, Rift Valley fever virus (RVFV), whose genome has three segments (S in 15 CPU hours / 7.1 minutes, M in 18 CPU hours / 10.3 minutes, L in 593 CPU hours, 553 minutes). Fourthly, we demonstrate the scalability of Panalyze by applying it to 2,358 mpox genomes (23 CPU hours, 249 minutes).

**Table 1.** Properties of selected virus PVGs. The Scaffold refers to the average scaffold size, which was from GFAstats such that the total length refers to the sum of the sequence lengths of the nodes in the PVG from ODGI. FMDV A stands for FMDV serotype A. FMDV C stands for FMDV serotype C. FMDV O stands for FMDV serotype O. Openness was estimated using the alpha values from Panacus.

| Virus | Region | Samples | Scaffold (bp) | Length (bp) | Nodes | Edges | Openness |
| --- | --- | --- | --- | --- | --- | --- | --- |
| LSDV | Genome | 121 | 150,524 | 156,966 | 7,203 | 9,827 | 1.88 |
| LSDV 5' end | 7.5 Kb | 132 | 7,500 | 8,954 | 790 | 1,037 | 1.80 |
| LSDV 3' end | 5 Kb | 132 | 5,001 | 7,198 | 443 | 572 | 1.64 |
| GTPV | Genome | 6 | 150,350 | 163,590 | 7,523 | 10,235 | 1.92 |
| SPPV | Genome | 31 | 149,834 | 151,742 | 3,136 | 4,354 | 1.04 |
| FMDV A | Genome | 142 | 8,028 | 15,793 | 13,102 | 21,235 | 0.81 |
| MPOX | Genome | 2,358 | 197,051 | 280,783 | 32,387 | 46,230 | 0.60 |
| PRCV | Genome | 15 | 27,660 | 32,086 | 7,559 | 10,233 | 1.13 |
| FMDV O | Genome | 441 | 8,178 | 16,879 | 14,064 | 22,993 | 0.99 |
| FMDV C | Genome | 18 | 8,148 | 10,779 | 5,959 | 8,359 | 1.14 |
| RVFV | S segment | 414 | 1,690 | 2,739 | 1,738 | 2,495 | 0.97 |
| RVFV | M segment | 302 | 3,885 | 6,182 | 3,639 | 5,091 | 0.93 |
| RVFV | L segment | 306 | 6,399 | 9,092 | 5,528 | 7,677 | 0.99 |

We repeated the analyses of the same sets of viral genomes using the nf-core/pangenome pipeline (Heumos et al 2024) for comparison with Panalyze using the same sets of viral genomes. The nf-core/pangenome pipeline version tested setup, environment and implementation was cloned locally and required at least 12 nodes.

Metrics were extracted for the pruned PVGs to ensure consistency of comparisons. Panalyze and the nf-core/pangenome pipeline provided identical results and the the nf-core/pangenome pipeline was faster overall. However, the RVFV segments S and M were too small to be assessed by nf-core/pangenome. In addition, Panalyze performs a wide range of PVG analysis and visualisations absent in nf-core/pangenome.

To run Panalyze on laptop environments, we created two instances of virtual machines on laptops. The high-resource instance had four cores with 8 GB RAM and the low-resource one had a single core with 4 GB RAM. We used a test dataset of 800 bp from the RVFV S segment for eight genomes.

## 3. Results

We created PVGs for large DNA (LSDV, mpox, GTPV, SPPV), small ssRNA (PRCV, FMDV) and segmented RNA (RVFV) viruses using Panalyze to illustrate its broad applicability to different virus types, as well as levels of diversity, sequence sizes and sample sizes (Table 1). The varied relative rate of nodes and edges compared to the sample sizes and average genome lengths illustrate how different the levels of sequence variability were. To illustrate the unusual patterns of variation in certain viral genomes, we extracted a portion of the LSDV PVG at which there are 15 mutations spanning two genes (Figure 1E). Phylogenetic reconstruction shows divergent haplotypes: selecting a single representative reference sequence for this region would bias downstream analyses.

We modelled the rate of discovery of new mutations as a function of the number of samples added to the PVGs, estimated with Panacus calculation of alpha. We observed that RVFV’s segments had a slightly open PVG, whereas PRCV and the FMDV serotypes had slightly closed ones, which contrasted with the closed LSDV PVG and the open mpox PVG (Table 1). Similar values were observed with Panacus. Our estimate of gamma for LSDV’s PVG (Panacus 0.04) was comparable to a previous estimate of 0.05 derived from a gene-based pangenome (Xie et al 2024).

## 4. Discussion

We present Panalyze, a scalable, portable, adaptable pipeline for virus PVG construction, analysis and annotation. It runs in NextFlow (Di Tommaso et al 2017) inside Docker (Merkel 2014) containers and is designed in a modular fashion. This allows Nextflow’s process management to improve thread allocation and accelerate task completion. Panalyze is written in domain-specific language (DSL) 2 and wraps a range of existing PVG tools to interpret PVGs rapidly to support exploration and optimisation. The selection of tools to run can be specified using a configuration file. Panalyze-generated PVGs uses PGGB and then runs a series of tasks to interpret this PVG using numerical, visual, graphical and other summaries for users to examine. This automation facilitates deeper engagement with complex datasets. Notably, it can analyse genomic regions of any size. Additionally, it can run on a scalable number of threads (from one to many), allowing it to be run locally on a laptop as well as on a HPC server. We demonstrate here how Panalyze can be applied to diverse specimens, including large DNA viruses, small ssRNA viruses, and segmented ones.

Although there are emerging tools for PVG construction, analysis and annotation for smaller genomes, no tool has yet bridged these nor automated them in Nextflow. Panalyze addresses this in a user-oriented reproducible manner. We hope that Panalyze can help the virus research community progress from using linear reference genomes to more accurate PVGs generated from more representative datasets. Moreover, we expect that superior PVG methods will underpin better novel or recombinant sample characterisation as well as routine PVG analysis and mutation detection. Consequently, Panalyze supports virus evolutionary and epidemiological genomics.

## Acknowledgements

We acknowledge comments and ideas from Dr. Caroline Wright (Pirbright Institute) and Dr. Lidia Dykes (Pirbright Institute).

