## Supplementary material for "Panalyze: automated virus pangenome variation graph construction, analysis and annotation": Suppl_data

**Supplementary Methods**

Panalyze is written in domain-specific language (DSL) 2 syntax and packaged in NextFlow v24.10.2 in a modular format. Each step or analysis has its own module that can be linked to subworkflow. We use NextFlow so that these processes could be imported easily into other subworkflows or pipelines. Portability is supported by packaging analysis dependencies in archived containers so that the local compute environment is the same in different HPC servers. We offer detailed documentation and community support through GitHub. We used Docker v27 and NextFlow v24.10.2 for each run that was implemented on a local HPC. The latter had 16 cores, 32 threads with 20 GB RAM and 100 GB scratch space. Each NextFlow run was allocated up to 40 threads.

As an example of the functional effects of this variation, we visualised eight nodes spanning 143 bp of a F4L tyrosine phosphatase gene at which there were three SNPs whose variation can be explored in a range of methods with VG (Garrison et al 2018), ODGI (Guarracino et al 2022), Bandage (Wick et al 2015) and GFAVIZ (Figure S1).

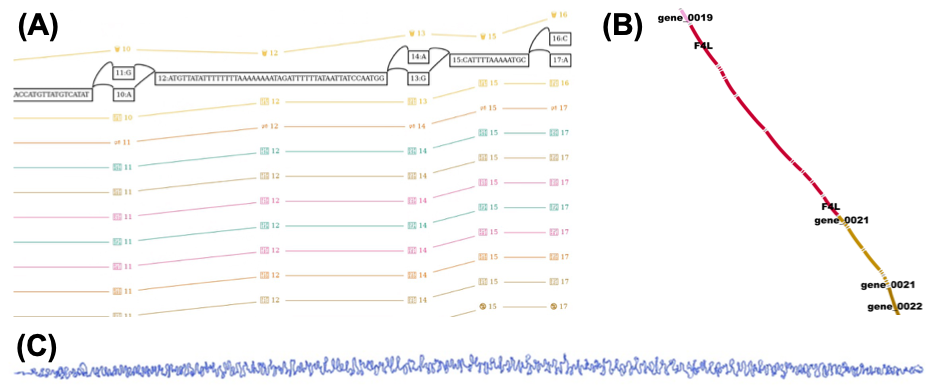

**Figure S1**. An example of PVG structure using LSDV: (A) Nodes labelled 9-17 linked to node 444 in gene encoding the F4L tyrosine phosphatase are shown, including SNPs at nodes 10/11, 13/14 and 16/17. The coloured lines represent different paths. (B) Bandage visualisation of the F4L gene and adjacent genes. (C) A visualisation of the whole LSDV PVG using GFAVIZ.
